# The shadowing effect of initial expectation on learning asymmetry

**DOI:** 10.1101/2022.11.22.517473

**Authors:** Jingwei Sun, Yinmei Ni, Jian Li

## Abstract

Evidence for positivity and optimism bias abounds in high-level belief updates. However, no consensus has been reached regarding whether learning asymmetries exists in more elementary forms of updates such as reinforcement learning (RL). In RL, the learning asymmetry concerns the sensitivity difference in incorporating positive and negative prediction errors (PE) into value estimation, namely the asymmetry of learning rates associated with positive and negative PEs. Although RL has been established as a canonical framework in interpreting agent and environment interactions, the direction of the learning rate asymmetry remains controversial. Here, we propose that part of the controversy stems from the fact that people may have different value expectations before entering the learning environment. Such default value expectation influences how PEs are calculated and consequently biases subjects’ choices. We test this hypothesis in two learning experiments with stable or varying reinforcement probabilities, across monetary gains, losses and gain-loss mixtures environments. Our results consistently support the model incorporating asymmetric learning rates and initial value expectation, highlighting the role of initial expectation in value update and choice preference. Further simulation and model parameter recovery analyses confirm the unique contribution of initial value expectation in accessing learning rate asymmetry.

**Author Summary:** While RL model has long been applied in modeling learning behavior, where value update stands in the core of the learning process, it remains controversial whether and how learning is biased when updating from positive and negative PEs. Here, through model comparison, simulation and recovery analyses, we show that accurate identification of learning asymmetry is contingent on taking into account of subjects’ default value expectation in both monetary gain and loss environments. Our results stress the importance of initial expectation specification, especially in studies investigating learning asymmetry.

## Introduction

When interacting with the uncertain environment, humans learn by trial-and-error, incorporating information into existing beliefs to accrue reward and avoid punishment, as reinforcement learning theory prescribes [1]. When an action leads to better-than-expected outcome and thus a positive prediction error is generated, such action tends to be repeated; in contrast, if an action is followed by a worse-than-expected outcome (negative prediction error), the tendency to repeat that action is reduced. Early reinforcement learning models typically assume that people’s sensitivities (learning rates) towards positive and negative prediction errors are the same[1–3]. Recently, however, evidence starts to emerge that the impacts of relatively positive and negative outcomes might be different[4–9], and distinct neural circuits may subserve learning from positive and negative prediction errors[10, 11].

Surprisingly, no consensus has been reached regarding the direction of learning asymmetry. In cases of high-level and ego-related belief updates, it has been shown that people tend to overestimate the likelihood of positive events and underestimate the likelihood of negative ones, a bias termed unrealistic optimism, possibly to maintain self-serving psychological status [12–16]. For example, when faced with new information about adverse life events, participants updated their beliefs more in response to desirable information (better than expected) than to undesirable information (worse than expected) [17–19] (but also see [20, 21]). However, results for the learning asymmetry in more elementary forms of updates such as reinforcement learning are rather mixed. While some studies using standard reinforcement learning paradigms have found that humans’ positive learning rates were larger than the negative ones, demonstrating an optimistic reinforcement learning bias [4, 22, 23]. Other studies, however, yielded opposite results with negative learning rates larger than the positive ones [6, 7, 24], consistent with the prevalent psychological phenomenon “bad is stronger than good” [25].

We hypothesize that part of the discrepancies in the previous literatures stems from the often less appreciated fact that the initial or default value expectation (*Q*_0_ in a Q-learning framework) plays a critical role in identifying the direction of learning asymmetry. In a standard two-arm bandit Q-learning paradigm, action value is updated by the product of learning rate (δ) and PE (δ), which is the difference between obtained reward (*R_t_*) and action value (*Q_t-1_*) of previous trial for specific trial *t*. Intuitively, setting the initial action value *Q*_0_ would have a direct impact on the calculation of immediate PE [26]. For example, if the endowed initial action value is lower than the true value per the action being selected, the positive prediction errors are up-scaled and negative ones down-scaled, creating an ostensible positivity bias (learning rate associated with positive PE is bigger than that of the negative PE). On the contrary, a negativity bias can emerge if the initial action value is mis-specified to be higher than the true value. However, a majority of recent studies focused on the role of learning rate in capturing participants’ behavior whereas considered *Q*_0_ as a mundane initialization parameter without a consensus as to how to initialize *Q*_0_. Indeed, while some recent studies set *Q*_0_ to zero, probably reflecting the fact that participants possess no information about options before entering the task [6–8, 23, 27, 28]; other studies adopted *Q*_0_ as the median or mean values of the possible option outcomes, corresponding to an *a priori* expectation of receiving different outcomes with equal probabilities [4, 28–30]. Few studies treated *Q*_0_ as a free parameter [31], due to the belief that the impact of initial expectation should be “washed out” after enough trials of learning.

However, it is plausible that there are significant individual differences in the initial expectation. Such initial expectation could reflect the internal motivation, or response vigor that participants carry into the task [32, 33]. In addition, the initial expectation might be susceptible to instructions or context cues, which have been shown to have clear impacts on participants’ choice behavior [31, 33–35]. Furthermore, contrary to the standard view, the initial value expectation may have long-lasting effects on subsequent choices due to the intricate interplay between choice selection and action value update. For example, if upfront interactions with a certain option widen the action value gap due to the specification of certain initial action values, then the lower valued option is less likely to be selected, making it harder to learn the true value of that option [6]. Therefore, RL models that do not take initial expectations into account may risk attributing variance in choice behavior to different causes, and also affect the estimation of the underlying learning rates.

To verify this hypothesis, we conducted two experiments where subjects were asked to select between probabilistically reinforced stimuli in the stable (Experiment 1) and random-walk (Experiment 2) probability environments. Two groups of subjects repeatedly chose from pairs of options with probabilistic binary reward outcomes to earn monetary rewards, avoid losses or both. We tested different variants of RL models against participants’ behavior with the focus on learning asymmetry and initial expectations. Our results showed that the RL model with asymmetric learning rates and individualized initial expectations performed best in both experiments 1 & 2. Further simulation and recovery analyses confirmed our results and demonstrated the characteristic impacts on learning asymmetry by omitting the initial expectation.

## Results

### Logistic regression and computational models

Twenty-eight subjects (one excluded due to technical problems) participated Experiment 1, where they were asked to choose from pairs of visual stimuli that were partially reinforced with fixed probabilities (Fig 1A). Experiment 1 consisted of two blocks (monetary gain and loss) and each block consisted of four pairs of options and their probabilities for winning (in Gain block) or losing (in Loss block) were 40-60%, 25-75%, 25-25% and 75-75%, respectively. Each pair of options was grouped into a mini-block and consisted of 32 trials.

**Figure 1.**
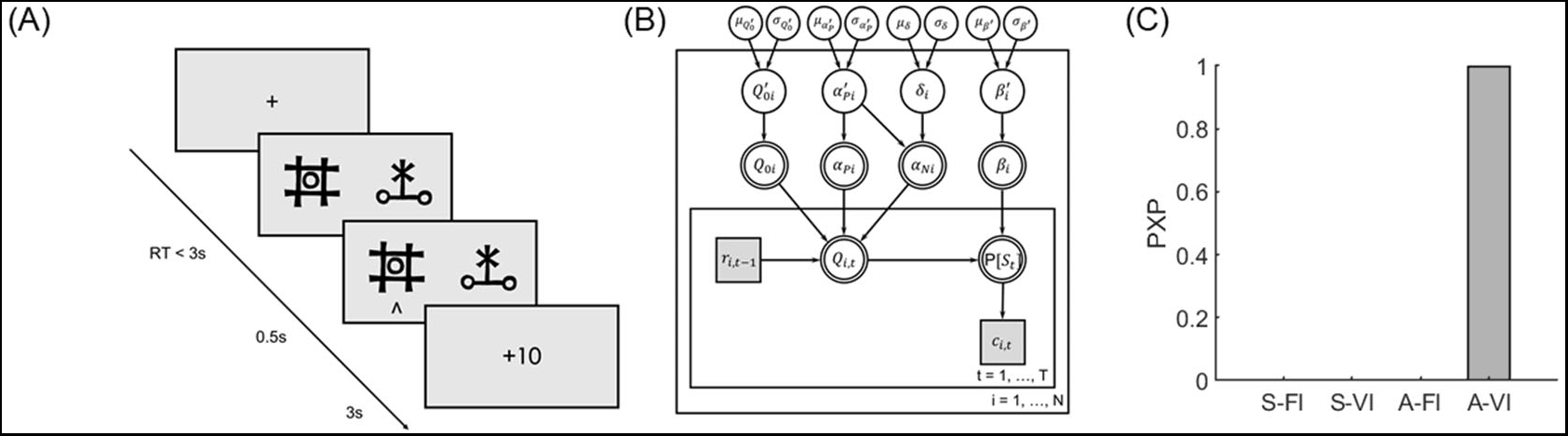
Experimental design and computational model of experiment 1 (stable probability). (A). Trial procedure of experiment 1. (B) Illustration of the hierarchical Bayesian modeling procedure. (C) Model comparison results.

Mixed-effect logistic regression (lme4 package in R v3.3.3 [36]) showed that subjects’ choices were sensitive to the past reward history (last trial outcome on stay probability: *β* = 0.958, *p* < 0.001), indicating that subjects did pay attention to the tasks and learned by trial-and-error. To test our hypothesis concerning learning asymmetry and initial expectation, we fitted the data with a standard Q-learning model assuming different learning rates for positive and negative prediction errors with individual initial expectation (A-VI). We also fitted three variants of this model, one with fixed initial expectation (A-FI, the initial expectation was 0.5 in gain, −0.5 in loss and 0 in mix condition), one with symmetric learning rates and initial expectation (S-VI), and lastly the one with fixed initial expectation and symmetric learning rates (S-FI). Deviance information criterion (DIC) analysis and Bayesian model selection indicated that the A-VI model performed the best in explaining subjects’ behavior with the protected exceedance probability (PXP) for the A-VI model at 99.9% (Fig 1C).

### Learning asymmetry revealed by the inclusion of initial expectation

As most of the previous literatures investigating learning asymmetry did not consider that initial expectation may vary across subjects, we specifically examined the difference of learning rates estimated from the A-VI and A-FI models. We found the direction of learning asymmetry suggested by these two models were different. While the positive learning rates appeared to be larger than the negative learning rates according to the A-FI model in both gain and loss conditions (Fig 2A, though not statistically significant, *p* = 0.265 for gain and *p* = 0.506 for loss, paired t-test), consistent with the positivity hypothesis [4, 22, 23], such pattern reversed course by incorporating initial expectation variation (A-VI model) in both the gain (Fig 2B, *p* < 0.001, paired t-test) and the loss condition (Fig 2B, *p* < 0.001, paired t-test). Importantly, there was no significant Pearson correlation between learning rates and initial expectation (*Q*_0_) in either gain or loss condition (in the best model, A-VI model), confirming the unique contribution of *Q*_0_ in explaining participants’ learning behavior (*r* = −0.120, *p* = 0.550 between *Q*_0_ & positive learning rate: *α*_P_; *r* = 0.235, *p* = 0.237 between *Q*_0_ & negative learning rate: *α*_N_ in the gain condition; *r* = 0.017, *p* = 0.935 between *Q*_0_ & *α*_P_, *r* = 0.362, *p* = 0.064 between *Q*_0_ & *α*_N_, in the loss condition).

**Figure 2.**
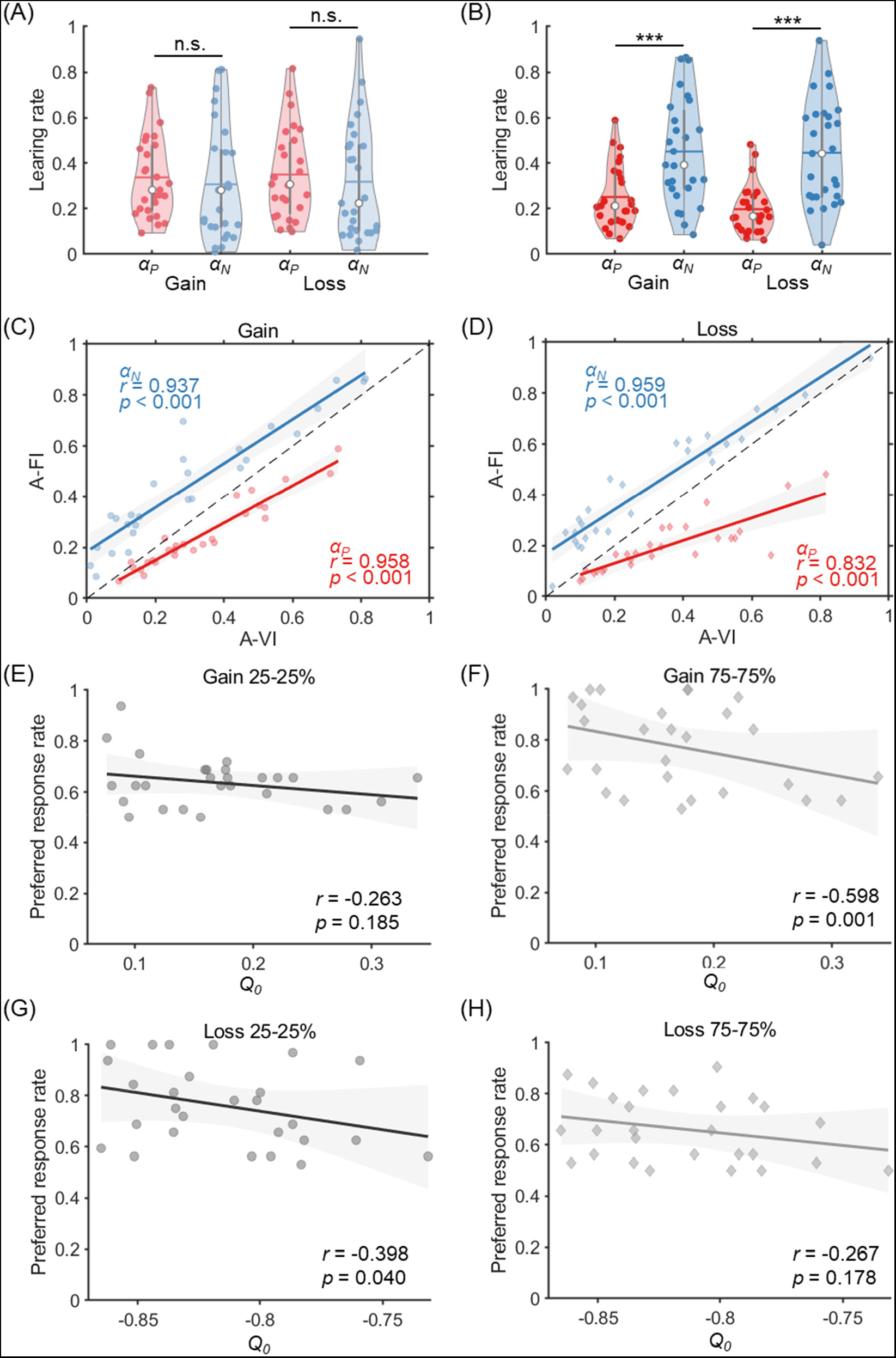
Model results of experiment 1. (A-B). Learning rates for gain and loss conditions estimated by the A-FI (A) and A-VI models(B). (C-D). Learning rate correlations between A-FI and A-VI models in the gain (C) and loss (D) conditions. (E-F). The correlation between preferred response rate (PRR) and *Q*_0_ (from A-VI model) in the gain 25-25% (E) and gain 75-75% (F) blocks. (G-H). Correlations of *Q*_0_ and PRR in the loss 25-25% (G) and loss 75-75% (H) blocks.

Despite the learning asymmetry reversal by considering individual *Q*_0_ in the A-VI model, however, closer examination of the learning rates estimated from the A-VI and A-FI models showed interesting correlation. Indeed, *α*_P_ and *α*_N_ were strongly correlated with their counterparts between the two models both for gain (*α*_P_: *r* = 0.958, *p* < 0.001; *α*_N_: *r* = 0.937, *p* < 0.001; Fig 2C) and loss conditions (*α*_P_: *r* = 0.832, *p* < 0.001; *α*_N_: *r* = 0.959, *p* < 0.001; Fig 2D), suggesting the relative rank of the individual difference in learning rates (positive or negative) is well preserved in both A-VI and A-FI models.

In experiment 1, we also included 25-25% and 75-75% blocks which according to previous literature might provide crucial evidence to support the optimistic reinforcement learning hypothesis [26, 28, 37]. We also tested such hypothesis and found that the ‘preferred response’ rate (PRR), a term defined as the choice rate of the option most frequently chosen by the subject and potentially reflects the tendency to overestimate certain option value, was correlated with *Q*_0_. More specifically, PRR was only negatively correlated with *Q*_0_ in the 75-75% gain condition (*r* = −0.598, *p* = 0.001; Fig 2F) and 25-25% loss condition (*r* = −0.398, *p* = 0.04; Fig 2G) where there was considerable mismatch between participants’ mean *Q*_0_ (mean *Q*_0_ = 0.170 and −0.815 in the gain and loss conditions) and the true action value (0.75 in the 75-75% gain condition and −0.25 in the 25-25% loss condition, respectively), indicating that PRR might instead be driven by the rather inaccurate initial expectation. Indeed, when the initial expectation was close to the true option value (25-25% gain condition and 75-75% loss condition), such correlation was not observed (Fig 2E, *r* = −0.263, *p* = 0.185 in the 25-25% gain condition; Fig 2H, *r* = −0.267, *p* = 0.178 in the 75-75% loss condition). These results suggest that as the discrepancy between individual and true *Q*_0_ grows larger, participants are more likely to experience extreme PEs and hence stick with an option that in fact has no obvious advantage.

### Model simulation and parameter recovery

To comprehensively investigate the influence of initial expectation on the estimation of learning rates, we further performed a model simulation analysis. We systematically varied the levels of the initial expectation (*Q*_0_ = 0, 0.25, 0.5, 0.75, 1) as well as the asymmetry of the positive and negative learning rates ((*α*_P_, *α*_N_) = (0.2, 0.6), (0.3, 0.5), (0.4, 0.4), (0.5, 0.3), (0.6, 0.2)) to simulate datasets using the A-VI model. Each combination of parameters generated 30 datasets with each dataset consisted of 30 hypothetical subjects, resulting in 750 (25 × 30) datasets in total. We then applied the same model fitting procedure with A-VI and A-FI models to the simulated datasets. For the purpose of exposition, we only simulated the gain condition.

As expected, the parameters were well-recovered by the A-VI model for all the parameter combinations (Fig 3A-C). On the contrary, when fitting without considering initial expectation differences across subjects (A-FI, *Q*_0_ = 0.5), both the positive and negative learning rates showed a systematic deviation from their true underlying values (Fig 3D-E). More specifically, when *Q*_0_ < 0.5, the positive learning rates were overestimated and the negative learning rates underestimated; whereas the positive learning rates were underestimated and the negative learning rates overestimated when *Q*_0_ > 0.5. The reason for such biases is due to the fact that when the true *Q*_0_ deviates from the assumed *Q*_0_(0.5), prediction errors caused by the misspecification of initial expectation can only be absorbed by rescaling the learning rates. Further learning rate asymmetry analysis demonstrated this pattern: the learning rate asymmetry (*α*_P_ − *α*_N_) was over estimated when the true initial expectation *Q*_0_ < 0.5 and underestimated when *Q*_0_ > 0.5 (Fig 3F). Furthermore, asymmetric learning model with another typical assumption of initial value (*Q*_0_ = 0) was also fitted to the simulation data and again produced estimation biases (Supplementary Fig 2), with the learning rate asymmetry (*α*_*P*_ − *α*_*N*_) underestimated when the true *Q*_0_ > 0 (Supplementary Fig 2C).

**Fig 3.**
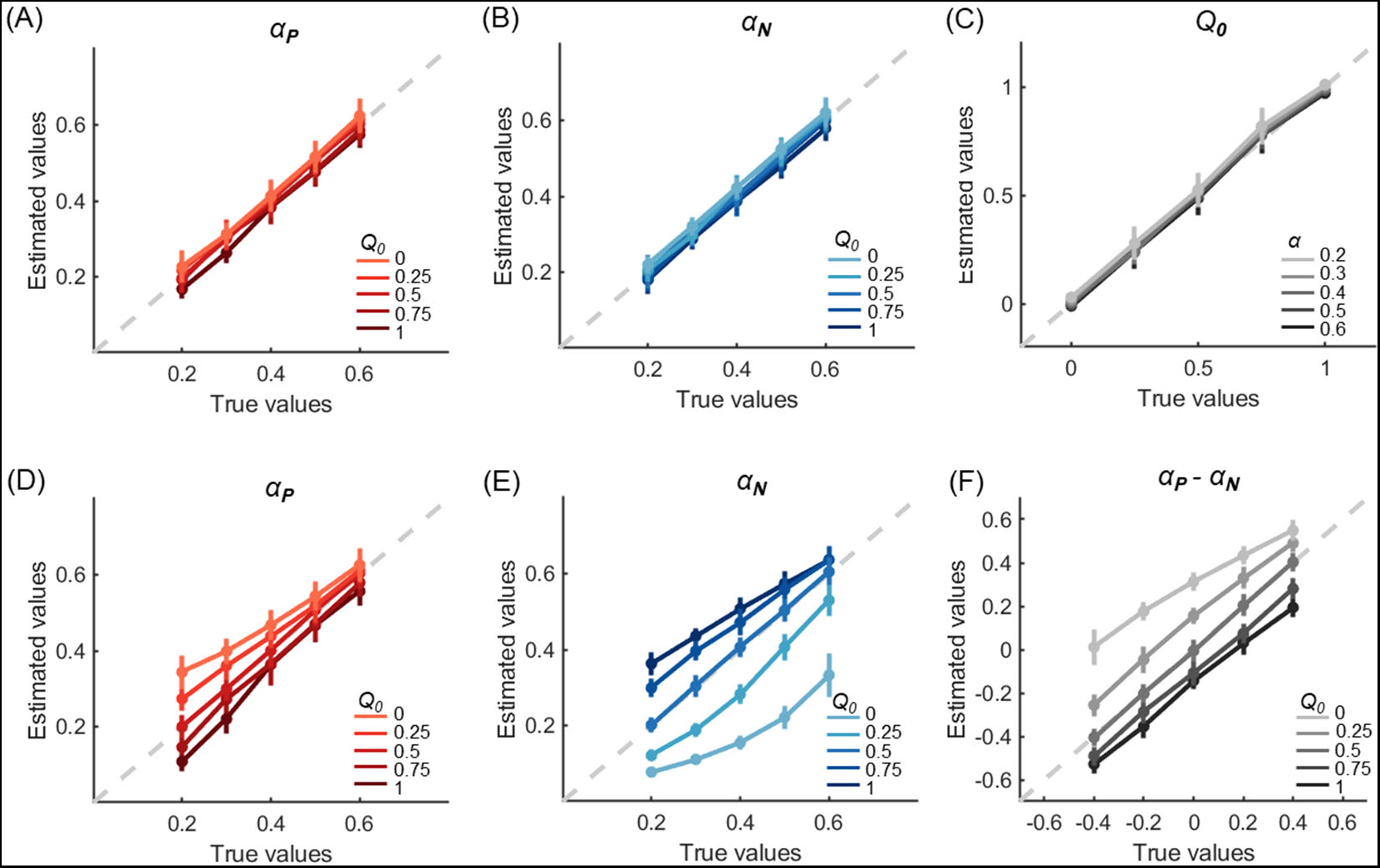
Simulation and parameter recovery for the Gain condition of experiment 1. Choice data were simulated using different combinations of positive/negative learning rates and initial expectations. Then, these data were fitted by the A-VI (A-C) and A-FI (D-F) models. The A-VI model faithfully retrieved the underlying parameters (A-C) whereas the A-FI model showed consistent deviation in parameter recovery (D-F). Error bars denote standard deviations across simulated subjects.

We also directly examined the estimated learning asymmetries with the posterior distribution of *μ_δ_*, the hyperparameter of the learning asymmetry in the A-VI and A-FI models for the simulated data (Fig 1b). For each combination of the underlying parameters, the estimated *μ_δ_* from the 30 datasets were pooled together to form the posterior distribution of *μ_δ_* (Fig 4). For the A-VI model, the learning asymmetry was correctly recovered for all initial expectation levels and learning rate pairs (Fig 4A). However, the learning asymmetry was only partially recovered for the A-FI model (Fig 4B, Supplementary Fig 3). Consistent with the learning rate estimation bias mentioned before, if *Q*_0_ < 0.5, the estimated positive learning rate tended to be larger than the negative learning rate (even if the true positive and negative learning rates were identical, or the true positive learning rate was smaller than the negative one) (Fig 4B red shaded areas). Likewise, if *Q*_0_ > 0.5, the estimated negative learning rate tended to be larger than the positive one (Fig 4B red shaded areas).

**Fig 4.**
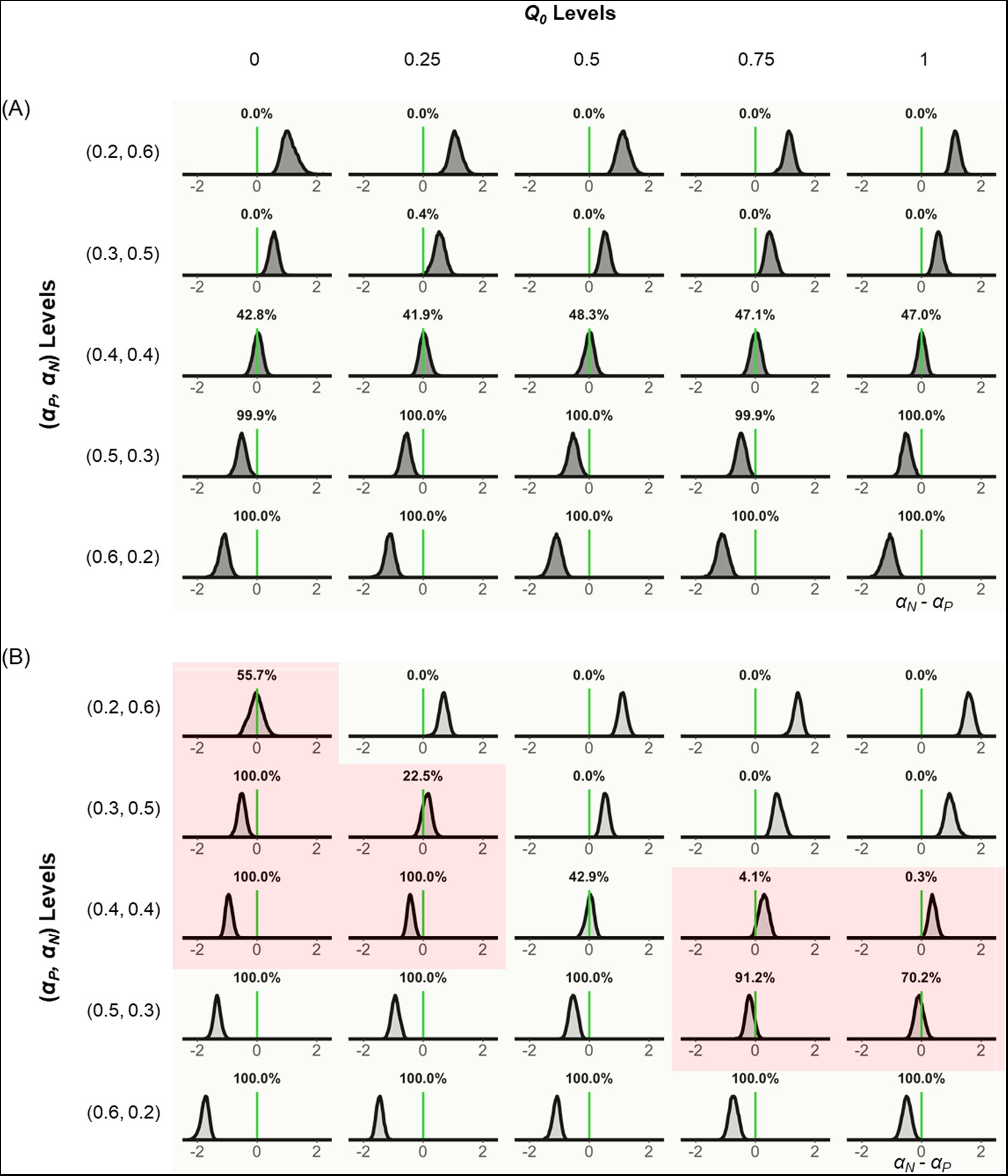
Recovered learning rate asymmetry for experiment 1 gain condition. The posterior distribution of *μ_δ_*, the hyper parameter of learning asymmetry for the A-VI model (A) and A-FI model (B). Light green in each distribution indicates faithful recovery (A-VI), whereas red shows the wrong categorization (A-FI).

### Generalization of the initial expectation effect to non-stable learning environment

To test the obstinate effect of initial expectation on learning behavior, we further collected participants’ choices in a non-stable learning environment (Experiment 2), where the reward (or punishment) probability of options gradually evolved over time (random walk with boundaries) and the learning sequence is longer than the stable environment (Fig 5A and 5B). In this experiment, we also included another condition of mixed valence options, where the outcome of an option is either positive (+10 points) or negative (−10 points). 30 subjects participated in this experiment. Similar model fitting procedure was applied, and the model comparison analysis found that the A-VI model outperformed the other three alternatives, with its protected exceedance probability larger than 99.9% (Fig 5C). Again, A-FI and A-VI models produced different learning rate asymmetry (Fig 5D-E). While A-FI model estimation only revealed significant learning asymmetry between positive and negative learning rates in the loss and mix conditions (*p* < 0.001 and *p* < 0.001 respectively, paired t-test) but not in the gain condition (*p* = 0.161; Fig 5D), the A-VI model showed consistent biased learning pattern across all three conditions, with the negative learning rate significantly larger than the positive learning rate (all *p*s < 0.001; Fig 5E). The learning rates revealed by these two models were also significantly correlated in all three conditions (Figs 5F-H; gain *α_P_*: *r* = 0.816, *p* < 0.001; gain *α_N_*: *r* = 0.916, *p* < 0.001; loss *α_P_*: *r* = 0.849, *p* < 0.001; loss *α_N_*: *r* = 0.828, *p* < 0.001; Mix *α_P_*: *r* = 0.900, *p* < 0.001; Mix *α_N_*: *r* = 0.919, *p* < 0.001). Similarly, we also ran model simulation and parameter recovery analysis for the gain trials in Experiment 2 (Fig 6), and the results confirmed that not specifying the initial expectation caused biased estimation of both the positive and negative learning rates: *α_P_* was overestimated and underestimated when *Q*_0_ was smaller or bigger than 0.5, respectively (Fig 6D). *α_N_*, however, was mainly underestimated (Fig 6E). The difference between *α_P_* and *α_N_* was mainly overestimated when *Q*_0_ < 0.5 and slightly underestimated when *Q*_0_ > 0.5 (Fig 6F). Finally, posterior distribution of *μ_δ_* in experiment 2 confirmed that learning asymmetry could be correctly identified at different *Q*_0_ levels when *Q*_0_ was treated as an individual parameter (Fig 7A), whereas mis-specification of learning difference would occur as a by-product of ignoring the heterogeneity of initial expectations (Fig 7B). Biased learning asymmetry was also induced when *Q*_0_ was fixed to be 0 in A-FI model recovery analysis (Supplementary Fig 3–4).

**Figure 5.**
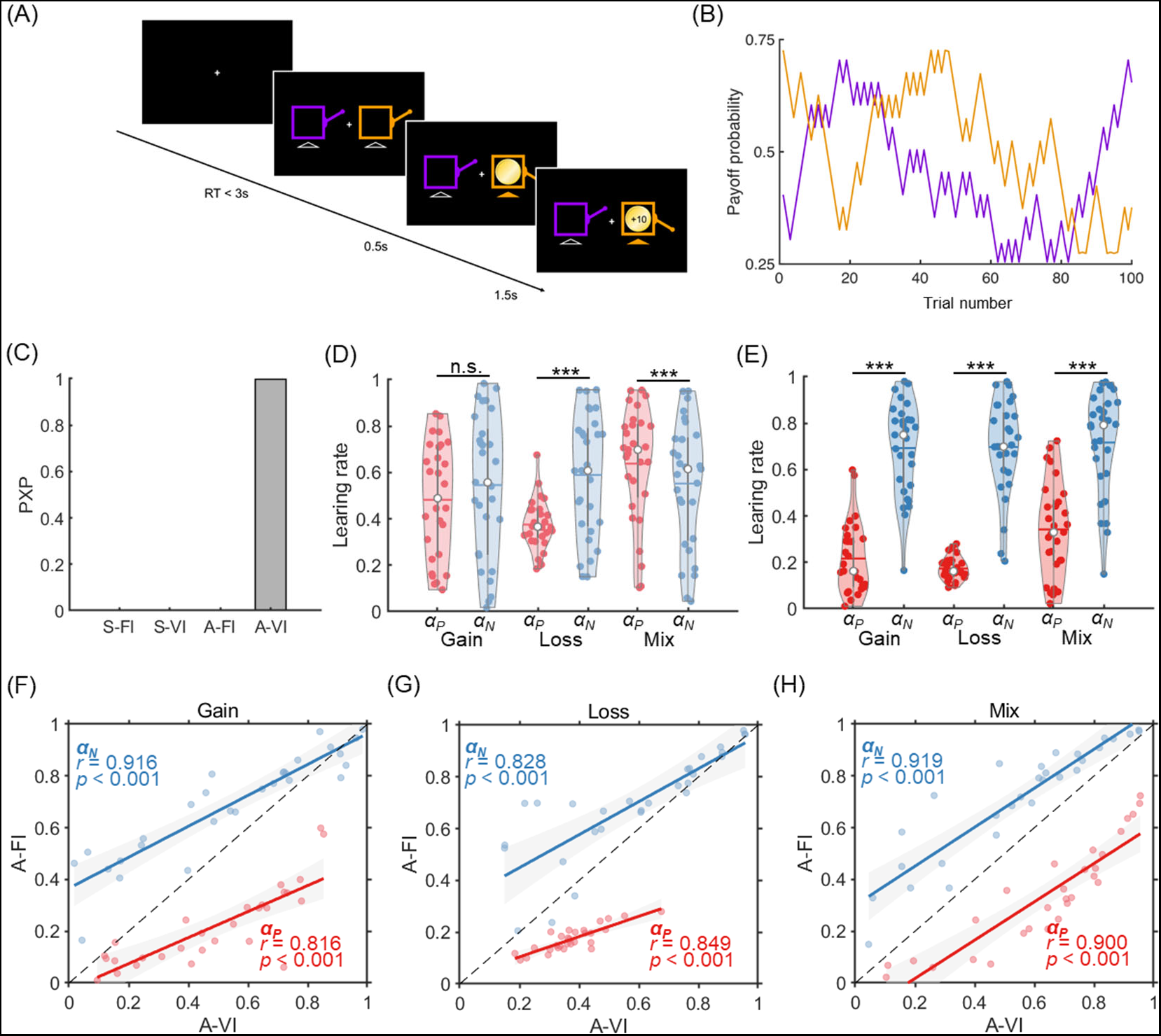
Experiment results of experiment 2. (A). A sample trial for experiment 2. (B). Example payoff probability sequences for the two slot machines (purple and orange). (C). Model comparison results for the 4 candidate models. (D-E). Consistent pattern of learning asymmetry was observed under the A-VI model for the gain, loss and mix conditions (E) but not for the A-FI (D) model. (F-H) Learning rates are positively correlated between A-FI and A-VI model estimation for all the gain (F), loss (G) and mix conditions (H).

**Figure 6.**
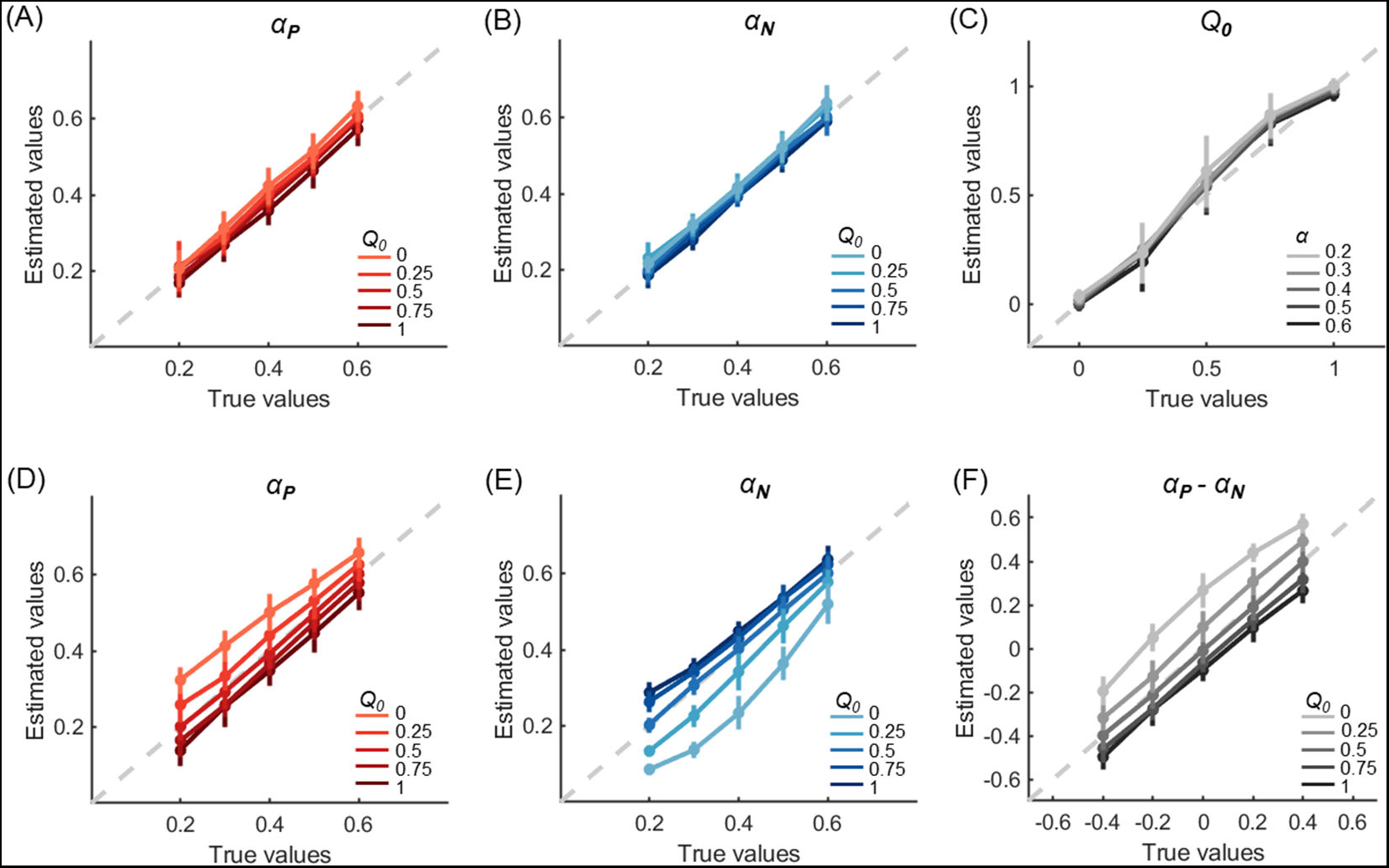
Simulation and parameter recovery for experiment 2 Gain condition. 1. Choice data were first simulated using different combinations of positive/negative learning rates and initial expectations and then submitted for model fitting and parameter recovery by the A-VI (A-C) and A-FI (D-F) models. The A-VI model faithfully retrieved the underlying parameters (A-C) whereas the A-FI model showed consistent deviation in parameter recovery (D-F). Error bars denote standard deviations across simulated subjects.

**Figure 7.**
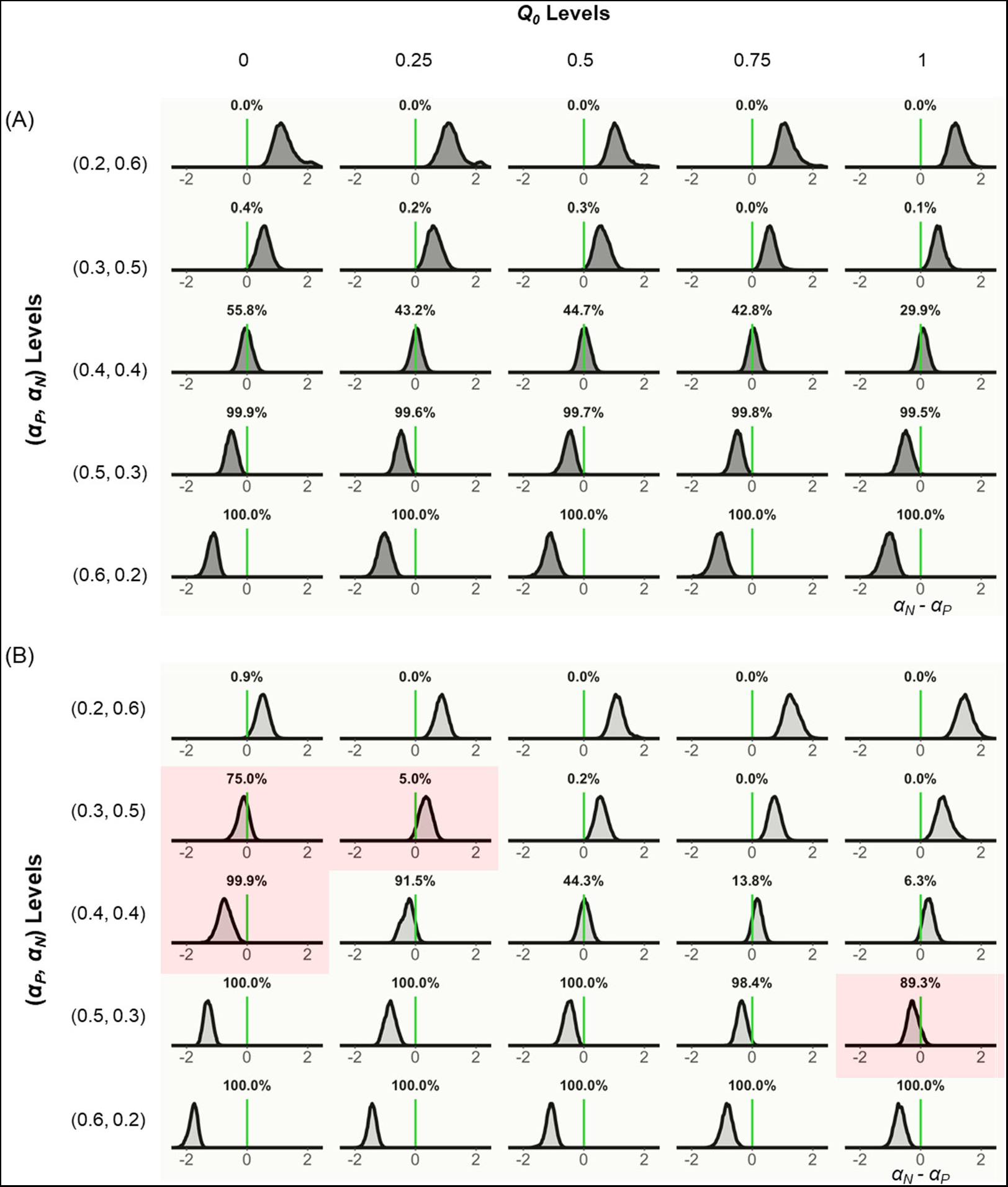
Recovered learning rate asymmetry for experiment 2 Gain condition. The posterior distribution of *μ_δ_*, the hyper parameter of learning asymmetry for the A-VI model (A) and A-FI model (B). Light green in each distribution indicates faithful recovery (A-VI), whereas red shows the wrong categorization (A-FI).

## Discussion

In two experiments, we tested and verified the hypothesis that the initial expectation has a profound impact on participants’ choice behavior, as opposed to the general assumption that so long as the trial numbers are long enough, the effect of initial expectation would be “washed out”. Interestingly, as a consequence, we also found that learning asymmetry (positive and negative learning rates) estimation can be consistently biased depending on the distance between the assumed and the true underlying initial expectation levels. We systematically tested these results in both stable (Experiment 1) and slowly evolving random-walk (Experiment 2) probabilistic reinforcement learning environments. For both experiments, the model with asymmetry learning rate and initial expectation parameters (A-VI) fitted subjects’ behavior best, suggesting the initial expectation parameter could capture additional variance of subjects’ behavior, above and beyond what can be explained by the learning asymmetry.

Previous literatures have linked state or action values to psychological mechanisms such as incentive salience, which maps “liked” objects or actions to “wanted” ones [33]. This line of research emphasized the critical role played by dopamine in assigning incentive salience to states or actions [38, 39]. Other research suggests that such value expectations also affect the strength or vigor of responding in free-operant behaviors [40], possibly with the evolvement of tonic dopamine. The motivational characteristic of action value suggests it is not only critical for generating PE, but also influencing how PE is obtained through choice selection. For example, when subjects were endowed with low expectations to start the gain task and received reward, the rather large positive PE would drive the selected option value up such that subjects tend to stick with this option and miss the opportunity to explore the other option. This is indeed what we observed in the equal probability conditions in experiment 1 (Fig. 2E-H): when subjects’ initial expectations (*Q*_0_) deviate from true option values, there were negative correlations between *Q*_0_ and the preferred response rates (Fig 2F&G); however, such correlation disappeared when *Q*_0_ was more consistent with option value (Fig 2E&H).

It is interesting to note that after removing the shadowing effects of initial expectation, results from both experiments revealed a consistent negativity bias in learning: people learn faster from negative PEs than from positive ones. This result holds across valence (gain and loss) and option reinforcement probability structures (stable and random-walk). Despite recent interests on learning asymmetries in belief, value and group impression updating [16, 17, 26, 37, 41], questions still remain regarding the direction and magnitude of the asymmetry. Although evidence starts to emerge to support a positivity bias (*α_P_* > *α_N_*) ranging from high-level belief update to more elementary forms of updates such as reinforcement learning [17, 26, 37], other studies seem to support a negativity bias (*α_P_* < *α_N_*) in learning [42–47]. One possibility to reconcile such discrepancy is by considering participants’ belief about the casual structure of the environment. For example, it has been shown that if the participants infer that experienced good (or bad) outcomes are due to a hidden cause, rather than the outcome distribution, they would learn relatively less from these outcomes, thus generating the putative negativity (or positivity) bias [16]. Here we propose another possibility: learning asymmetry estimation may be over-shadowed by participants’ initial expectation. Indeed, computational modeling analysis may yield learning asymmetry with different directions depending on the specification of default *Q*_0_, even when learning is symmetric (Fig 3F and Fig 6F).

It should also be noted that the relative rank of the individual difference in learning rates (positive or negative) is well preserved, with or without the consideration of initial expectations. In fact, correlation analyses of both the *α_P_* and *α_N_* from the A-FI and the A-VI models showed they were positively correlated across different conditions (Figs 2C-D; Fig 5F-H). However, when inferences are to be drawn about learning asymmetry, that is, the comparison of *α_P_* and *α_N_*, the effect of initial expectation starts to emerge. Previous literatures have shown that other factors such as response autocorrelation might also influence whether learning asymmetry can be identified and proposed model-free methods to mitigate estimation bias [48, 49]. Our current study adds to this line of research by demonstrating the necessity of including initial expectation level to better capture subjects’ learning behavior in different learning environments (stable and random-walk reinforcement probability), different outcome valences (gain, loss or mixed reward) and different lengths of learning sequences (short or long).

In summary, here we demonstrate that initial expectation level plays a significant role in identifying learning asymmetry in a variety of learning environments, supported by both computational modeling and model simulation and parameter recovery analyses. Our findings help pave the way for future studies about learning asymmetry, which has been implicated in a range of learning and decision making biases in both healthy people [15, 50–52], as well as those who suffer from psychiatric and neurological diseases [53, 54].

## Methods

### Ethics statement

The experiments had been approved by the Institutional Review Board of School of Psychological and Cognitive Sciences at Peking University. All subjects gave informed consent prior to the experiments.

### Subjects

The study consisted of two experiments. 28 subjects participated in Experiment 1 (14 female; mean age 22.3 ± 3.2), of which one participant (male) was excluded from analysis due to technical problems. 30 subjects participated in Experiment 2 (16 female; mean age 22.1 ± 2.4) and one participant (male) was excluded due to the exclusive selection of one-side option on the computer screen during the experiment (97%).

### Behavioral tasks

In each experiment, subjects performed a probabilistic instrumental learning task in which they chose between different pairs of visual cues to earn monetary rewards or avoid monetary losses. In Experiment 1, characters from the Agathodaemon alphabet were used as cues and their associative outcome probability were stationary. Outcome valence was manipulated in two blocks: in the Gain block, the possible outcomes for each cue were either gaining 10 points or zero, whereas in the Loss block, outcomes were either losing 10 points or zero. In each block there were four probability pairs of 40/60%, 25/75%, 25/25% and 75/75%, respectively. Probability conditions were grouped into mini-blocks, with 32 trials for each condition. There’s a minimum of 5 seconds’ rest between mini-blocks, and a minimum of 20 seconds’ rest between two blocks. The visual cues for each condition were randomly selected, and the assignment of probabilities to the cues were counterbalanced across conditions. Participants started with two practice mini-blocks (5 trials each) before the experiment using different visual cues and outcome probabilities. At the end of the experiment, points earned by the participants were converted to monetary payoff using a fixed ratio and participants earned ¥45 on average.

Within each block, a trial started with a fixation cross at the center of the computer screen (1 s), followed by the presentation of cue pairs (maximum 3 s), during which subjects were required to choose either the left or right cue by pressing the corresponding buttons on the keyboard. An arrow (0.5 s) appeared under the cue (Fig 1A) to indicate the chosen option immediately after subjects made their choices, followed by the outcome of that trial. If subjects responded faster than the 3s time limit, the remaining time was added to the duration of fixation presentation of next trial. If no choice was made within the 3s response time window, a text message “Please respond faster” was displayed for 1.5 s and subjects needed to complete the trial again to ensure 32 choice selections were collected for each pair of cues.

The task design of experiment 2 was similar to experiment 1, and subjects were required to choose between two slot machines. The major distinction of experiment 2 was that the outcome probabilities of the stimuli followed a random-walk scheme instead of remaining stable [31, 55]. At the beginning of the task, slot machine outcome probabilities were independently drawn from a uniform distribution with boundaries of [0.25, 0.75]. Following each trial, the probabilities were diffused either up or down, equiprobably and independently, by adding or subtracting 0.05. The updated probabilities were then reflected off the boundaries [0.25, 0.75] to maintain them within the range. We tested three types of outcome valence as Gain, Loss, and Mix (in which the possible outcomes were either earning 10 points or losing 10 points) blocks. Each block consisted of choosing from a pair of slot machines for 100 trials. The color of slot machines was randomly selected, and the order of the three blocks were counterbalanced.

### Computational models

The Q-learning algorithm has been used extensively to model subjects’ trial-by-trial behavior during learning [56–59]. It assumes subjects learn by updating the expected value (*Q* value) for each action based on the prediction error (*δ*). In our study, we allowed the learning rates for positive and negative prediction errors to be different. After every trial *t*, the value of the chosen option is updated as follows:

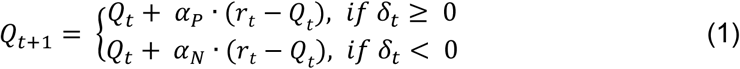

The term *r_t_* - *Q_t_* is the prediction error (*δ_t_*) in trial *t* and we set the reward, *R_t_* = −1, 0, 1 for losing, 0, and winning, respectively. *α_P_* and *α_N_* are the positive and negative learning rates and are constrained in the range of [0, 1]. The initial expectation for each option, *Q*_0_, is set as a free parameter, constrained in the range between the worst and the best outcome of that option. We assumed the initial expectation for all options were the same for each individual. We refer to this model as the asymmetric reinforcement learning model with variable initial expectation (A-VI).

The probability of choosing one option over the other is described by the softmax rule, with the inverse temperature *β* constrained in [0, 20]:

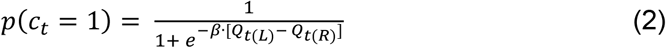

Here, *Q*_*t(L)*_ and *Q*_*t(R)*_ are the *Q* value for left and right options in trial *t*. We also considered other variant models of RL. The first one is A-FI, where the initial expectation *Q*_0_ were set at the mean outcome in the gain, loss and mix blocks (0.5, - 0.5 and 0) respectively, corresponding to an initial expectation of 50% chance of receiving either outcome. The second one is S-VI, where the learning rates for positive and negative prediction errors are the same (*α_P_* = *α_N_*). The last one is S-FI, where *Q*_0_s were set at the mean outcomes and with identical learning rates for positive and negative prediction errors. For the fixed initial expectation models (A-FI and S-FI), we also tested their performance with *Q*_0_ = 0 in the gain and loss conditions.

### Bayesian hierarchical modeling procedure and model comparison

We applied a Bayesian hierarchical modeling procedure to fit the models. In contrast to the traditional point estimate method, such as maximum likelihood, the Bayesian hierarchical method can estimate the posterior distribution of the parameters at the individual level as well as the group level in a mutually constraining fashion to provide more stable and reliable parameter estimation [60–62]. Take the example of A-VI model (Fig 1B), *r_i,t-1_* refers to the outcome received by subject *i* at trial *t*-1 and *c_i,t_* is the choice of subject *i* at trial *t*. The individual-level parameters were transformed using the Ф transformation, the cumulative density function of the standard normal distribution, to constrain the parameter values in their corresponding boundaries. In order to directly capture the effect of interest [62, 63], i.e. the learning rate asymmetry, we modeled the negative learning rate as the sum of the positive learning rate and the difference between negative and positive learning rates. Specifically, for each parameter *θ* (*θ* ∈ {*Q*_0_, *α_P_*, *β*}) with [*θ*_*min*_, *θ*_*max*_] as its boundary, *θ* = *θ*_*min*_ + Ф(*θ*^′^) x (*θ*_*max*_ - *θ*_*min*_). Parameters *θ*′ were drawn from hyper normal distributions with mean *μ*_θ_′ and standard deviation *σ*_θ_′. A normal prior was assigned to the hyper means *μ_θ′_* ~ *N*(0, 2) and a half-Cauchy prior to the hyper standard deviations *σ_θ_′* ~ *C*(0, 5). Negative learning rate was specified as 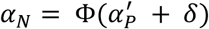, where *δ* was set the same way as *θ*′. The three alternative models were specified in a similar manner. Data from different outcome valence conditions was modeled separately.

Model fitting was performed using R (v3.3.3) and RStan (v2.17.2). For each model, 6000 samples were collected after a burn-in of 4000 samples on each of four chains, leading to a total of 24,000 samples collected for each parameter (representing the posterior distribution of the corresponding parameter). For each parameter, we computed a trimmed mean by discarding 10% samples from each side to obtain the robust estimation of the corresponding parameters [64].

Given the parameter samples, we computed deviance information criterion (DIC) for each model and used it to compare our candidate models’ performance [65]. We further calculated the protected exceedance probability (PXP), which indicates the probability that a specific model is the best model among the candidates, based on the group-level Bayesian model selection method [66, 67].

### Model simulations and parameter recovery

To test the robustness of our results, we performed a comprehensive parameter recovery analysis. For each task (stable or random-walk probability scheme), we generated hypothetical choices using the best performing model (A-VI model) with different initial expectation levels and different learning rates levels. We tested the gain condition parameter recovery for both experiment 1 (Fig 3 and Fig 4) and 2 (Fig 6 and Fig 7), respectively. Specifically, we considered five levels of initial expectation, where *Q*_0_ equals 0, 0.25, 0.5, 0.75 and 1, and five pairs of positive and negative learning rates, where (*α_P_*, *α_N_*) equals to (0.2, 0.6), (0.3, 0.5), (0.4, 0.4), (0.5, 0.3) and (0.6, 0.2). For each combination of the initial expectation and learning rates, we simulated 30 datasets, leading to a total of 750 (30 × 25 *Q*_0_ and learning rates combinations) datasets for each task. Each dataset consists of 30 hypothetical subjects. *β* was fixed to 5 for all datasets. For each dataset, we fitted models with and without parameterizing the initial expectation (where initial expectation was 0.5 or 0) using the same Bayesian model fitting method described above.

## Acknowledgements

JL is supported by the National Natural Science Foundation of China Grants (31871140, 32071090), National Science and Technology Innovation 2030 Major Program (No. 2021ZD0203702).

## Author contributions

Conceived and designed the experiments: J.S and J.L. Performed the experiments:

J.S. Analyzed the data: J.S and Y.N. Wrote the paper: J.S, Y.N and J.L.

---

**STable 1.** Model DICs. Model fitting results. Model 1, 3, 4 & 6 were reported in the main results. We also considered models where the …. was fixed at 0 instead of the mean outcome (model 2 & 5) for gain and loss conditions. Across two experiments, the A-VI model (M6) consistently performed better than all the other candidates.

| Model | Experiment1 | Experiment2 |
| --- | --- | --- |
| M1: S_FI<br>$Q_0$ is 0.5(gain), -0.5(loss), 0(mix) | 6442 | 7146 |
| M2: S_FI<br>$Q_0$ is 0(gain), 0(loss), 0(mix) | 7027 | 7233 |
| M3: S_VI<br>$Q_0$ is free parameter | 6122 | 6997 |
| M4: A_FI<br>$Q_0$ is 0.5(gain), -0.5(loss), 0(mix) | 6114 | 7066 |
| M5: A_FI<br>$Q_0$ is 0(gain), 0(loss), 0(mix) | 6926 | 7053 |
| M6: A_VI<br>$Q_0$ is free parameter | 6028 | 6811 |

**SFig 1.**
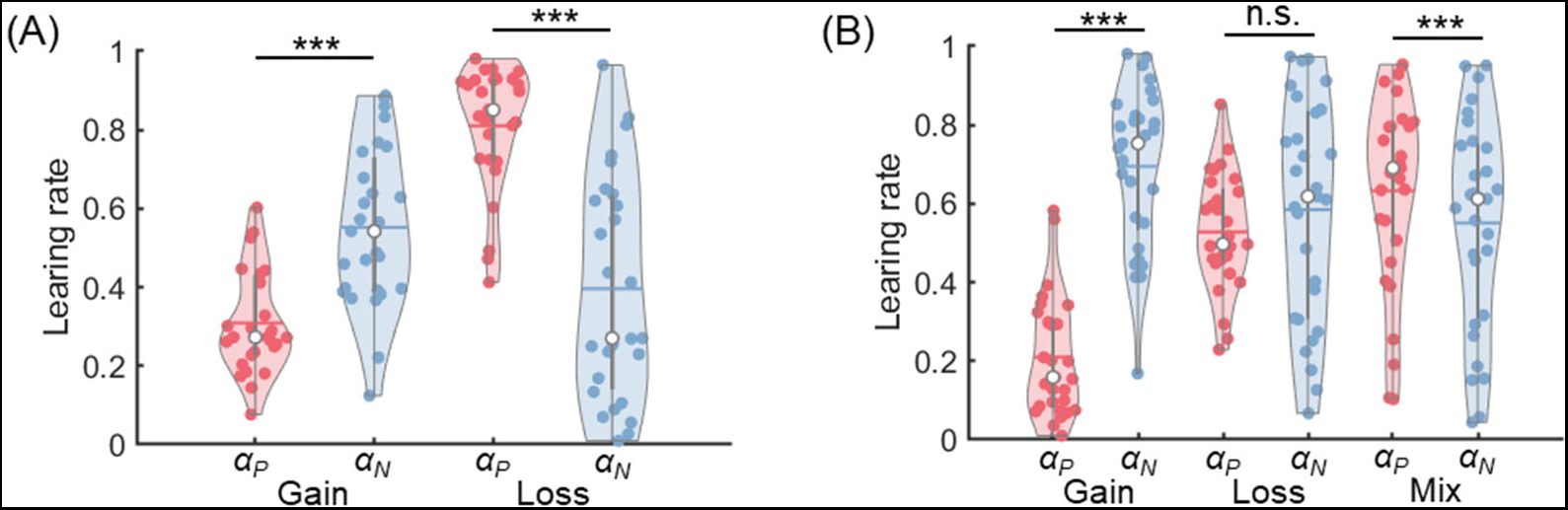
Learning rates estimated from Model 5 (M5) in two experiments. (A) In experiment 1, **α_P_** was significantly smaller than **α_N_** in the gain condition (paired t-test, *p* < 0.001) and larger in the loss condition (*p* < 0.001). (B) In experiment 2, **α_P_** was smaller and larger than **α_N_** in the gain and mix condition (*p_s_* < 0.001), respectively, and there was no significant difference between **α_P_** and **α_N_** in the loss condition (*p* = 0.145).

**SFig 2.**
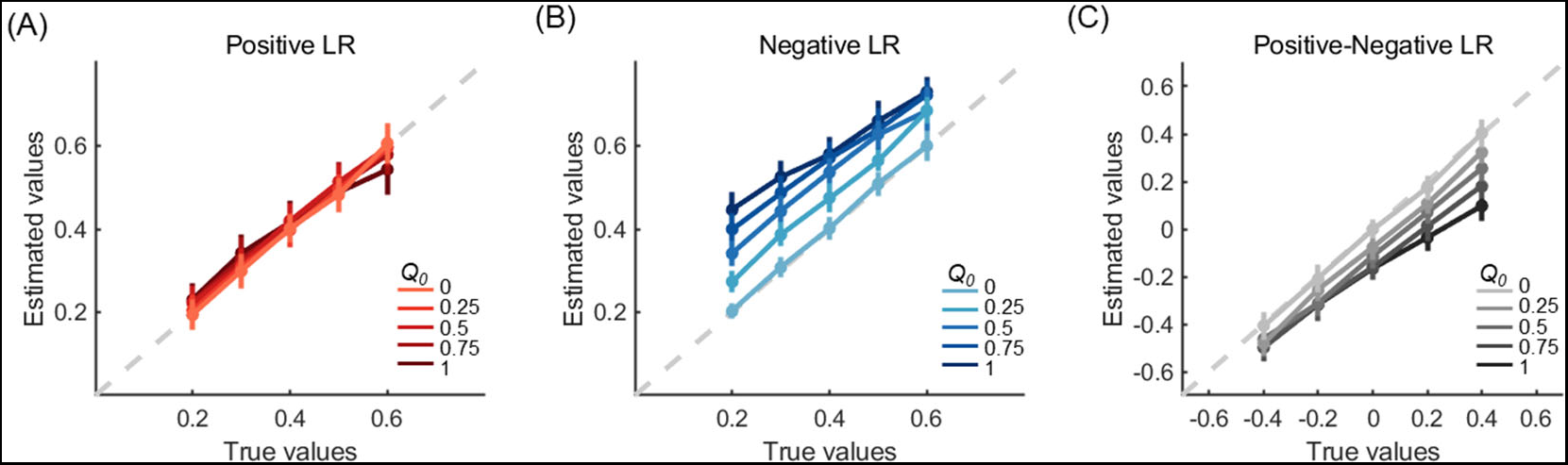
Simulation and parameter recovery for experiment 1 Gain condition. Choice data were simulated using different combinations of positive/negative learning rates and initial expectations and then fitted by the A-FI model with initial expectation *Q*_0_ = 0 (M5). Error bars denote standard deviations across simulated subjects.

**SFig 3.**
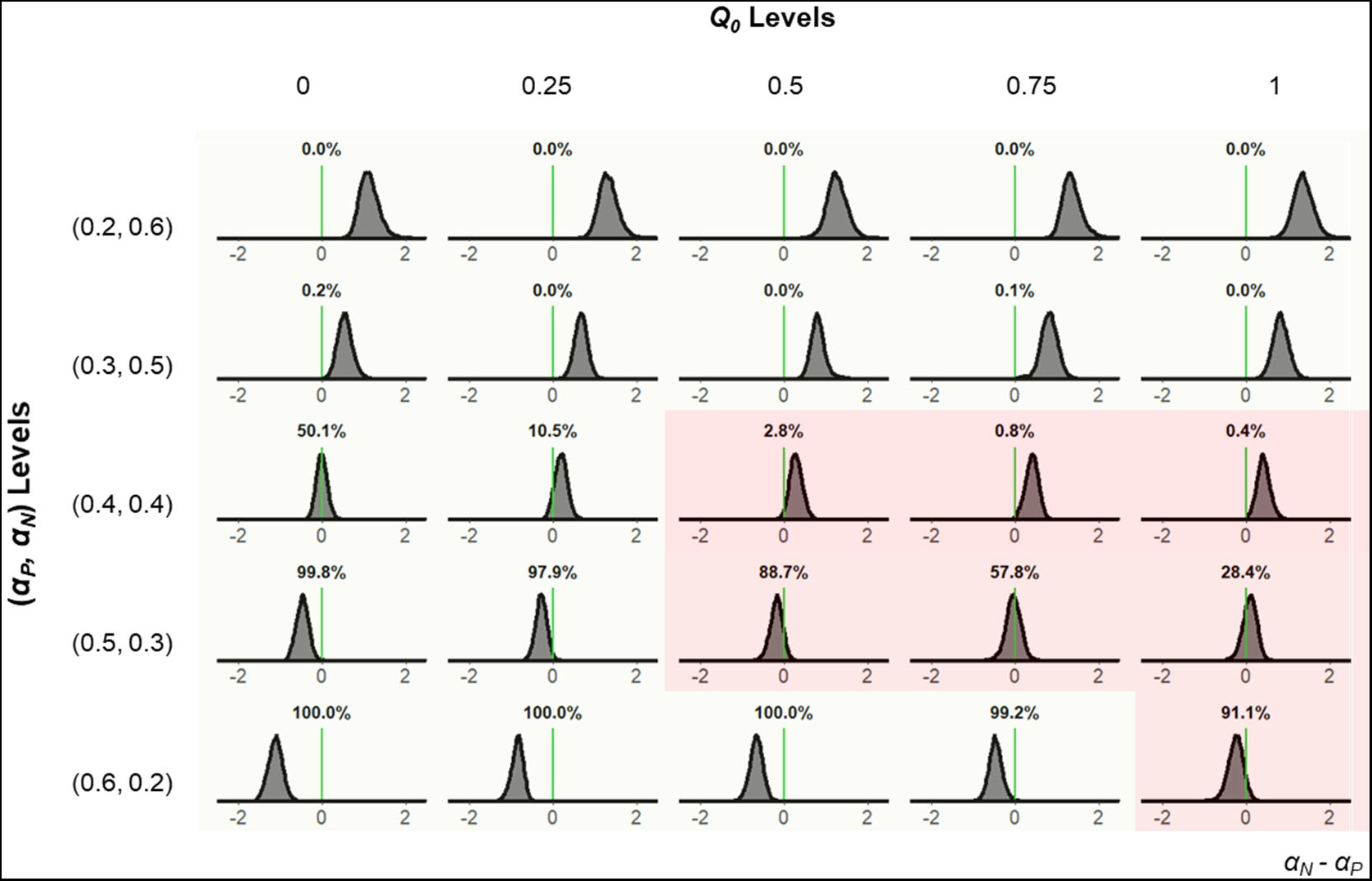
Recovered learning rate asymmetry for the gain condition of experiment 1. The posterior distribution of *μ_δ_*, the hyper parameter of learning asymmetry for the A-FI model with initial expectation *Q*_0_ = 0 (M5). Light green in each distribution indicates faithful recovery, whereas red shows the wrong categorization.

**SFigure 4.**
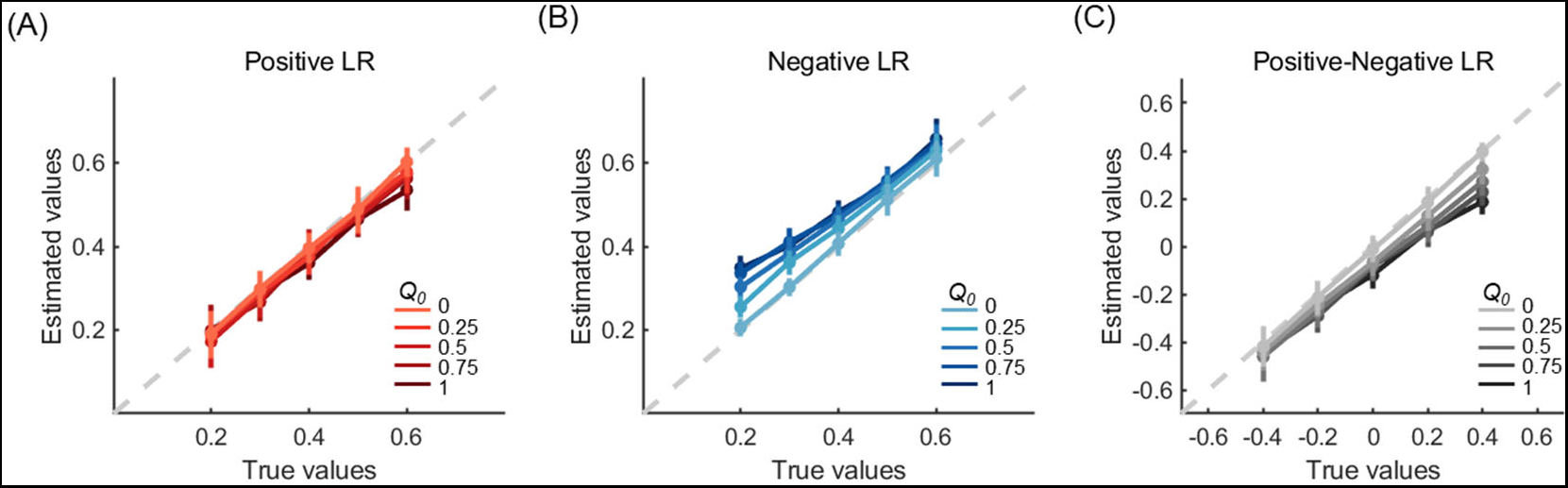
Simulation and parameter recovery for the gain condition of experiment 2. Choice data were simulated using different combinations of positive/negative learning rates and initial expectations and then fitted by the A-FI model with initial expectation *Q*_0_= 0 (M5). Error bars denote standard deviations across simulated subjects.

**SFig 5.**
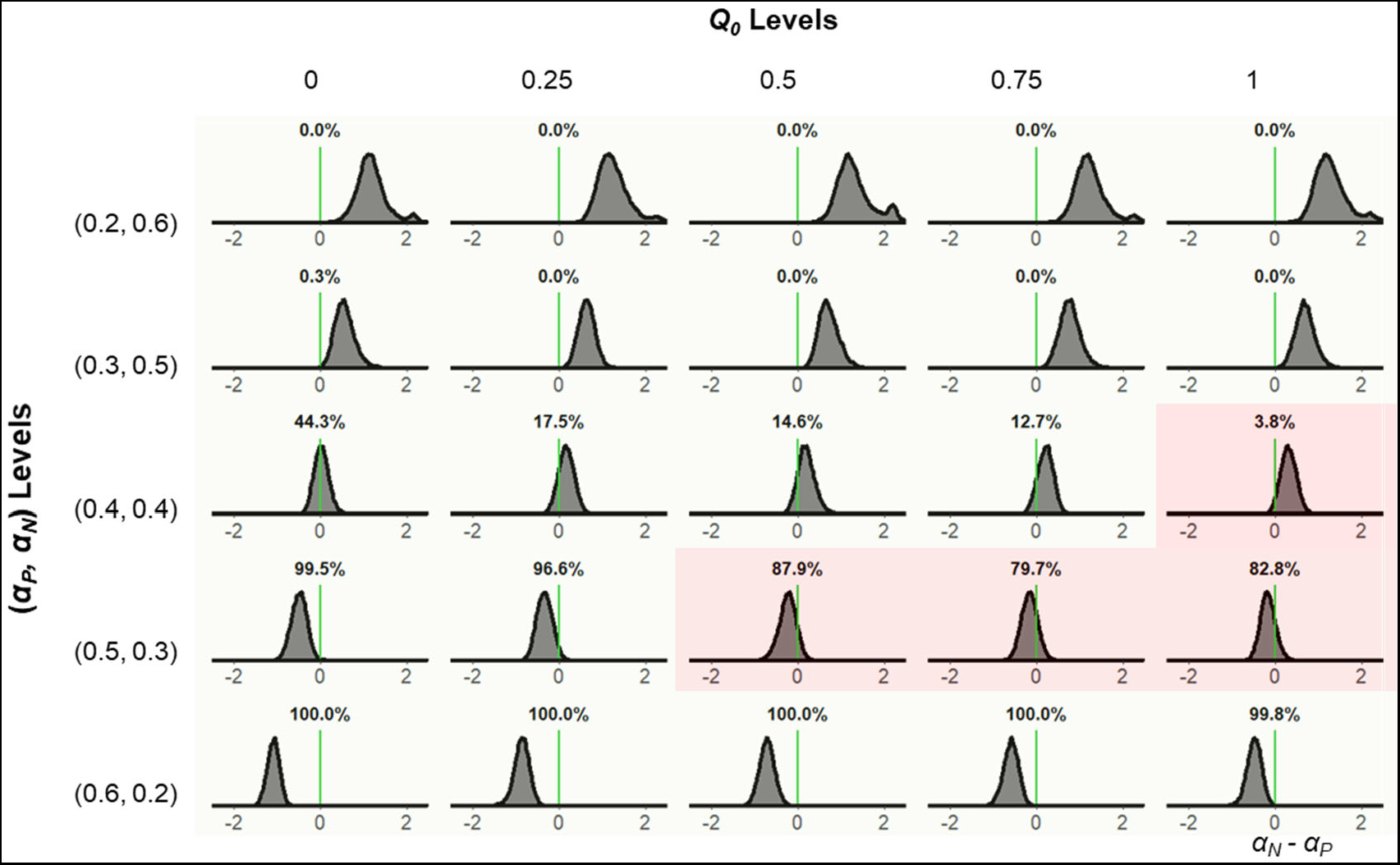
Recovered learning rate asymmetry for the gain condition in experiment 2. The posterior distribution of **μ_δ_**, the hyper parameter of learning asymmetry for the A-FI model with initial expectation *Q*_0_ = 0 (M5). Light green in each distribution indicates faithful recovery, whereas red shows the wrong categorization.

## References

1. Sutton RS, Barto AG. Reinforcement learning: An introduction: MIT press; 1998.

2. Pessiglione M, Seymour B, Flandin G, Dolan RJ, Frith CD. Dopamine-dependent prediction errors underpin reward-seeking behaviour in humans. Nature. 2006;442(7106):1042.

3. O’Doherty JP, Hampton A, Kim H. Model-based fMRI and its application to reward learning and decision making. Ann N Y Acad Sci. 2007;1104:35–53.

4. Lefebvre G, Lebreton M, Meyniel F, Bourgeois-Gironde S, Palminteri S. Behavioural and neural characterization of optimistic reinforcement learning. Nature Human Behaviour. 2017;1(4):0067.

5. Sharot T, Korn CW, Dolan RJ. How unrealistic optimism is maintained in the face of reality. Nature neuroscience. 2011;14(11):1475–9.

6. Niv Y, Edlund JA, Dayan P, O’Doherty JP. Neural prediction errors reveal a risk-sensitive reinforcement-learning process in the human brain. Journal of Neuroscience. 2012;32(2):551–62.

7. Gershman SJ. Do learning rates adapt to the distribution of rewards? Psychonomic bulletin & review. 2015;22(5):1320–7.

8. Frank MJ, Doll BB, Oas-Terpstra J, Moreno F. Prefrontal and striatal dopaminergic genes predict individual differences in exploration and exploitation. Nature Neuroscience. 2009;12(8):1062–8.

9. Frank MJ, Moustafa AA, Haughey HM, Curran T, Hutchison KE. Genetic triple dissociation reveals multiple roles for dopamine in reinforcement learning. Proceedings of the National Academy of Sciences. 2007;104(41):16311–6.

10. Frank MJ, Seeberger LC, O’Reilly RC. By carrot or by stick: Cognitive reinforcement learning in Parkinsonism. Science. 2004;306(5703):1940–3.

11. Kravitz AV, Tye LD, Kreitzer AC. Distinct roles for direct and indirect pathway striatal neurons in reinforcement. Nature neuroscience. 2012;15(6):816.

12. Weinstein ND. Unrealistic optimism about future life events. Journal of personality and social psychology. 1980;39(5):806–20.

13. Spiegelhalter DJ, Best NG, Carlin BP, Van Der Linde A. Bayesian measures of model complexity and fit. Journal of the Royal Statistical Society Series B, Statistical methodology. 2002;64(4):583–639.

14. Eil D, Rao JM. The Good News-Bad News Effect: Asymmetric Processing of Objective Information about Yourself. American Economic Journal: Microeconomics. 2011;3(2):114–38.

15. Sharot T, Garrett N. Forming Beliefs: Why Valence Matters. Trends Cogn Sci. 2016;20(1):25–33.

16. Dorfman HM, Bhui R, Hughes BL, Gershman SJ. Causal Inference About Good and Bad Outcomes. Psychol Sci. 2019;30(4):516–25.

17. Sharot T, Korn CW, Dolan RJ. How unrealistic optimism is maintained in the face of reality. Nat Neurosci. 2011;14(11):1475–9.

18. Sharot T, Guitart-Masip M, Korn CW, Chowdhury R, Dolan RJ. How dopamine enhances an optimism bias in humans. Curr Biol. 2012;22(16):1477–81.

19. Bromberg-Martin ES, Sharot T. The Value of Beliefs. Neuron. 2020;106(4):561–5.

20. Shah P, Harris AJ, Bird G, Catmur C, Hahn U. A pessimistic view of optimistic belief updating. Cogn Psychol. 2016;90:71–127.

21. Garrett N, Sharot T. Optimistic update bias holds firm: Three tests of robustness following Shah et al. Conscious Cogn. 2017;50:12–22.

22. Dorfman HM, Bhui R, Hughes BL, Gershman SJ. Causal Inference About Good and Bad Outcomes. Psychological science. 2019;30(4):516–25.

23. Ting C-C, Palminteri S, Lebreton M, Engelmann JB. The Elusive Effects of Incidental Anxiety on Reinforcement-Learning. Journal of experimental psychology Learning, memory, and cognition. 2021;48(5):619–42.

24. Christakou A, Gershman SJ, Niv Y, Simmons A, Brammer M, Rubia K. Neural and psychological maturation of decision-making in adolescence and young adulthood. Journal of cognitive neuroscience. 2013;25(11):1807–23.

25. Baumeister RF, Bratslavsky E, Finkenauer C, Vohs KD. Bad is stronger than good. Review of general psychology. 2001;5(4):323–70.

26. Palminteri S, Lebreton M. The computational roots of positivity and confirmation biases in reinforcement learning. Trends Cogn Sci. 2022;26(7):607–21.

27. Palminteri S, Justo D, Jauffret C, Pavlicek B, Dauta A, Delmaire C, et al. Critical Roles for Anterior Insula and Dorsal Striatum in Punishment-Based Avoidance Learning. Neuron. 2012;76(5):998–1009.

28. Palminteri S, Lefebvre G, Kilford EJ, Blakemore SJ. Confirmation bias in human reinforcement learning: Evidence from counterfactual feedback processing. PLoS Comput Biol. 2017;13(8):e1005684.

29. Bornstein AM, Khaw MW, Shohamy D, Daw ND. Reminders of past choices bias decisions for reward in humans. Nature Communications. 2017;8:15958.

30. Van Slooten JC, Jahfari S, Knapen T, Theeuwes J. How pupil responses track value-based decision-making during and after reinforcement learning. Plos Computational Biology. 2018;14(11):25.

31. Li J, Daw ND. Signals in human striatum are appropriate for policy update rather than value prediction. The Journal of neuroscience. 2011;31(14):5504–11.

32. Niv Y, Daw ND, Joel D, Dayan P. Tonic dopamine: opportunity costs and the control of response vigor. Psychopharmacology. 2007;191(3):507–20.

33. McClure SM, Daw ND, Montague PR. A computational substrate for incentive salience. Trends Neurosci. 2003;26(8):423–8.

34. Doll BB, Jacobs WJ, Sanfey AG, Frank MJ. Instructional control of reinforcement learning: a behavioral and neurocomputational investigation. Brain Res. 2009;1299:74–94.

35. Palminteri S, Khamassi M, Joffily M, Coricelli G. Contextual modulation of value signals in reward and punishment learning. Nat Commun. 2015;6:8096.

36. Bates D, Machler M, Bolker BM, Walker SC. Fitting Linear Mixed-Effects Models Using lme4. Journal of Statistical Software. 2015;67(1):1–48.

37. Lefebvre G, Lebreton M, Meyniel F, Bourgeois-Gironde S, Palminteri S. Behavioural and neural characterization of optimistic reinforcement learning. Nature human behaviour. 2017;1(4).

38. Berridge KC, Robinson TE. What is the role of dopamine in reward: hedonic impact, reward learning, or incentive salience? Brain research Brain research reviews. 1998;28(3):309–69.

39. Ikemoto S, Panksepp J. The role of nucleus accumbens dopamine in motivated behavior: a unifying interpretation with special reference to reward-seeking. Brain research Brain research reviews. 1999;31(1):6–41.

40. Niv Y, Daw ND, Joel D, Dayan P. Tonic dopamine: opportunity costs and the control of response vigor: Dopamine - revisited. Psychopharmacologia. 2007;191(3):507–20.

41. Burke CJ, Tobler PN, Baddeley M, Schultz W. Neural mechanisms of observational learning. Proc Natl Acad Sci U S A. 2010;107(32):14431–6.

42. Christakou A, Gershman SJ, Niv Y, Simmons A, Brammer M, Rubia K. Neural and psychological maturation of decision-making in adolescence and young adulthood. J Cogn Neurosci. 2013;25(11):1807–23.

43. Gershman SJ. Do learning rates adapt to the distribution of rewards? Psychon Bull Rev. 2015;22(5):1320–7.

44. Niv Y, Edlund JA, Dayan P, O’Doherty JP. Neural prediction errors reveal a risk-sensitive reinforcement-learning process in the human brain. J Neurosci. 2012;32(2):551–62.

45. Pulcu E, Browning M. Affective bias as a rational response to the statistics of rewards and punishments. eLife. 2017;6.

46. Wise T, Dolan RJ. Associations between aversive learning processes and transdiagnostic psychiatric symptoms in a general population sample. Nature communications. 2020;11(1):4179-.

47. Wise T, Michely J, Dayan P, Dolan RJ. A computational account of threat-related attentional bias. PLoS computational biology. 2019;15(10):e1007341–e.

48. Seymour B, Daw ND, Roiser JP, Dayan P, Dolan R. Serotonin selectively modulates reward value in human decision-making. J Neurosci. 2012;32(17):5833–42.

49. Katahira K. The statistical structures of reinforcement learning with asymmetric value updates. Journal of mathematical psychology. 2018;87:31–45.

50. Bénabou R, Tirole J. Mindful Economics: The Production, Consumption, and Value of Beliefs. The Journal of economic perspectives. 2016;30(3):141–64.

51. Bénabou R, Tirole J. Self-Confidence and Personal Motivation. The Quarterly journal of economics. 2002;117(3):871–915.

52. Sharot T, Rollwage M, Sunstein CR, Fleming SM. Why and When Beliefs Change. Perspectives on Psychological Science. 2022:17456916221082967.

53. Maia TV, Frank MJ. From reinforcement learning models to psychiatric and neurological disorders. Nature neuroscience. 2011;14(2):154.

54. Maia TV, Conceicao VA. The Roles of Phasic and Tonic Dopamine in Tic Learning and Expression. Biol Psychiatry. 2017;82(6):401–12.

55. Daw ND, Gershman SJ, Seymour B, Dayan P, Dolan RJ. Model-based influences on humans’ choices and striatal prediction errors. Neuron. 2011;69(6):1204–15.

56. Sutton RS, Barto AG. Reinforcement learning: An introduction: MIT press; 2018.

57. Dayan P, Abbott L. Theoretical neuroscience: computational and mathematical modeling of neural systems. Journal of Cognitive Neuroscience. 2003;15(1):154–5.

58. Jahfari S, Ridderinkhof KR, Collins AG, Knapen T, Waldorp LJ, Frank MJ. Cross-task contributions of frontobasal ganglia circuitry in response inhibition and conflict-induced slowing. Cerebral Cortex. 2018;29(5):1969–83.

59. Daw ND, O’Doherty JP, Dayan P, Seymour B, Dolan RJ. Cortical substrates for exploratory decisions in humans. Nature. 2006;441(7095):876.

60. Ahn W-Y, Haines N, Zhang L. Revealing neurocomputational mechanisms of reinforcement learning and decision-making with the hBayesDM package. Computational Psychiatry. 2017;1:24–57.

61. Ahn W-Y, Krawitz A, Kim W, Busemeyer JR, Brown JW. A model-based fMRI analysis with hierarchical Bayesian parameter estimation. 2013.

62. Sokol-Hessner P, Raio CM, Gottesman SP, Lackovic SF, Phelps EA. Acute stress does not affect risky monetary decision-making. Neurobiology of stress. 2016;5:19–25.

63. McCoy B, Jahfari S, Engels G, Knapen T, Theeuwes J. Dopaminergic medication reduces striatal sensitivity to negative outcomes in Parkinson’s disease. Brain. 2019;142(11):3605–20.

64. Acerbi L, Vijayakumar S, Wolpert DM. On the origins of suboptimality in human probabilistic inference. PLoS computational biology. 2014;10(6):e1003661.

65. Spiegelhalter DJ, Best NG, Carlin BP, Van Der Linde A. Bayesian measures of model complexity and fit. Journal of the Royal Statistical Society: Series B (Statistical Methodology). 2002;64(4):583–639.

66. Stephan KE, Penny WD, Daunizeau J, Moran RJ, Friston KJ. Bayesian model selection for group studies. Neuroimage. 2009;46(4):1004–17.

67. Rigoux L, Stephan KE, Friston KJ, Daunizeau J. Bayesian model selection for group studies—revisited. Neuroimage. 2014;84:971–85.

